# Effect of straw incorporation and nitrification inhibitor on nitrous oxide emission in various cropland soils and its microbial mechanism

**DOI:** 10.1101/2021.05.26.445903

**Authors:** Ya-Bo Zhang, Feng Liu, Jun-Tao Wang, Hang-Wei Hu, Ji-Zheng He, Li-Mei Zhang

**Author notes:** For correspondence: Jun-Tao Wang, or.

## Abstract

Nitrification inhibitor and straw incorporation are widely used to improve crop nitrogen use efficiency in agricultural soil, but their effects on nitrous oxide (N_2_O) emission across different soil types and the microbial mechanisms remain less understood. In this study, we used controlled experiment and DNA-based molecular analysis to study how nitrification inhibitor (dicyandiamide, DCD) and straw incorporation affect soil nitrogen balance, N_2_O emission and microbial nitrifiers/denitrifers in three distinct agricultural soils (the black, fluvo-aquic and red soils) across China. Both DCD and straw incorporation improved nitrogen balance by increasing NH_4_^+^ and decreasing NO_3_^-^ in all soils. DCD tended to decrease N_2_O emission from all soils especially the fluvo-aquic one, while straw incorporation reduced N_2_O emission only in the fluvo-aquic soil but increased N_2_O emission in the other two especially the red soil (by ∼600%). T-RFLP analysis revealed that the denitriers community structure are distinct among the three soils but was not strongly affected by DCD or straw incorporation. qPCR analysis revealed that DCD or straw incorporation had no significant effect on nitrifier abundance but increased nitrous oxide reductase *nosZ* gene abundance in the black/fluvo-aquic soil rather than the red soil. Structural equational modelling further confirmed that, when accounting for treatments and soil properties, *nosZ* gene abundance is the only biological factor significantly determined N_2_O emission in different soil types. Taken together, our work advanced the knowledge on the agricultural practices and N_2_O emission in cropland soils, suggesting that straw incorporation may not be a good choice for the red and black soil areas; management practices should be used as *per* soil type to balance between nitrogen use efficiency and N_2_O emission.

## Introduction

As a powerful greenhouse gas, nitrous oxide (N_2_O) is 256 times stronger than carbon dioxide in warming potential, and is widely recognized as a major contributor to ozone depletion (IPCC, 2014; Ravishankara et al., 2009). The use of synthetic nitrogen fertilizers in agricultural production is the biggest source of N_2_O emissions. It was estimated that 60% of the global N_2_O emission (about 4.1 Tg N yr^−1^) could be attributed to synthetic nitrogen fertilizers application (c.a. 140 Tg N yr^−1^) (IPCC, 2013). In the context of the global food crisis and climate change, it is particularly urgent to optimize fertilization strategies to increase nitrogen fertilizer use efficiency (NUE) while reducing N_2_O emissions from agricultural soils.

Straw incorporation (SI) and nitrification inhibitor (NI) are common practices employed to improve the NUE in cropland soils (Di et al., 2014), but their performances on regulating soil N_2_O emissions remained uncertain. For example, straw incorporation potentially increased the N_2_O emission in an aquic cropland soil (Xu et al., 2019), but may not affect the N_2_O emissions in a red soil under traditional synthetic fertilizer treatment (Wu et al., 2020). The widely used inhibitor dicyandiamide (DCD), even though largely suppressed the N_2_O emissions in an acidic loamy clay soil (Ning et al., 2018), might slightly increase the N_2_O emission from agricultural soils when globally assessed (Lam et al., 2017). These results indicated that the influence of SI and NI on N_2_O emission can be largely limited by soil conditions especially soil types, and a mechanistic interpretation on the N_2_O emission pattern across different soil types is essential for deploying fertilization strategies to mitigate N_2_O emissions and improve crop NUE.

In cropland soils, nitrous oxide is mostly emitted as an intermediate product of heterotrophic denitrification (NO_3_^−^→NO_2_^−^→NO^-^→N_2_O→N_2_) or/and as a by-product of ammonia oxidation (Butterbach-Bahl et al., 2013; Hu et al., 2015). The former pathway is driven by denitrifiers containing nitrite reductase (*nirK*) and/or nitrous oxide reductase (*nosZ*) genes. The latter is driven by ammonia-oxidizing bacteria/archaea (AOB/AOA) that encode ammonia-monooxygenase α-subunit gene (*amoA*). Changes of these functional microbial groups are essential for mechanistic interpretation on the response N_2_O emission to fertilization strategies (Chen et al. 2013). Straw incorporation doubled the abundance of *nosZ* gene, the only known biological link that reduces N_2_O emissions, in an acidic soil (Miller et al. 2012), but potentially increase the emissions of N_2_O from cropland soils across large spatial scales (Zhao et al. 2020). Previous study showed that DCD stressed paddy soil N_2_O emission via inhibiting the metabolic activity of AOB rather than AOA (Zhou et al. 2020). DCD was also reported to potentially reduce denitrification activity via reducing denitrification substrate (NO_3_^-^) concentration rather than the abundance of *nirK* gene in a paddy soil (Meng et al., 2020). Considering that soil nitrifier and denitrifer communities are highly variable across environmental gradient (Jones and Hallin 2010, Hu et al. 2015), how DCD and straw incorporation affect the nitrifier/denitrifier and regulate the N_2_O emissions across different soil types remains uncertain.

The black, fluvo-aquic, and red soils are representative of the main part of the agricultural areas across eastern China that accounts for more than half of the crop productions in China. Proper agricultural management policy across these areas is necessary for crop safe and environmental sustainability. Here, we used a controlled experiment to determine the performance of SI and NI on N_2_O in these three distinct soils. The responses of denitrifying and ammonia-oxidizing microorganisms were characterized by DNA molecular analysis, such as quantitative PCR and Terminal Restriction Fragment Length Polymorphism (T-RFLP). We aimed (1) to compare the performance of SI and NI on N_2_O emissions in different soil types, and (2) to unravel the microbial mechanism by which NI and SI affect soil N_2_O emissions among different soils. We wish our work can provide data support for deploying suitable fertilization strategies to both improve the crop NUE and mitigate soil N_2_O emissions in different agricultural areas.

## Materials and methods

### Soil basic information and controlled experiment setup

The soils used in the pot experiment were collected from agricultural areas in Taoyuan (TY, 111.26°E, 28.55°N), Xuchang (XC, 113. 48°E, 34. 08°N) and Gongzhuling (GZL, 124.49°E, 43.32°N), respectively. According to the World Reference Base (WRB), they were classified as Red soil (pH 4.43), Fluvo-aquic soil (pH 7.18) and Black soil (pH 5.74), respectively. The average annual temperature and precipitation in TY, GZL, and XC sites were 16.5°C/1440mm, 5.6°C/615mm and 13.9°C/595mm, respectively. Crop residuals and stones were removed, screened through a 2-mm sieve before transported to the greenhouse.

The pot experiment was set up in May, 2015. For each soil type, there were four fertilization treatments including unfertilized control (CK), using nitrogen fertilizer only, i.e., using urea equivalent to 200 kg N ha^−1^ (N), addition of 5% nitrification inhibitor (N+NI) and incorporation of maize straw (N+SI) with nitrogen fertilizer. Each treatment had four replicates. The corn straw used in N+SI treatment was collected from the same sites where the soils were collected. Phosphorus and potassium (equivalent to 90 kg P_2_O_5_ ha^−1^ and 90 kg K_2_O ha^−1^, respectively) were applied as the basal fertilizers for all the treatments. All pots (with 10 kg soil each) were irrigated to 60% water hold capacity (WHC) before seeding. The pots (Φ: 25cm × H: 25cm) were randomly arranged in greenhouse with constant temperature at ∼20°C. A widely grown maize variety *Zhengdan 985* was used in this experiment.

### Gas sampling and N_2_O measurement

Soil N_2_O flux were monitored using the static chamber/gas chromatograph as described before (Ding et al., 2015). Briefly, the soil column was covered by a Φ: 25cm × H: 60 cm transparent plexiglass chamber (with electric fan installed on the top to mix the air in the headspace) and a water-filled groove was used to seal the system. The sample collection time was from 10:00 to 11:00 on the 2, 4, 6, 8, 10, 12, 16, 20, 27 days after seeding. There were three time points (at 0, 15th, 30th min) for each collection; at each time point, a syringe is used to collect gas samples (30 ml) from the chamber. These samples were stored in glass cylinders and then determined by a gas chromatography (Agilent 7890B, Santa Clara, CA, USA) with electron capture detector (µECD). In all, 1,296 gas samples were collected and measured. Calculation of N_2_O flux was performed via a linear regression model (Wang et al., 2018b),

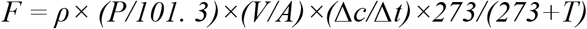

where *F* indicates N_2_O flux (µg N_2_O-N m^-2^ h^−1^), *ρ* indicates N_2_O density at 0°C and 101.3 KPa (kg m^-3^), *V* indicates chamber volume (m^3^), *A* indicates chamber surface area, *Δc/Δt* indicates N_2_O accumulation rate within chamber (ppbv N_2_O-N h^−1^), *T* (Celsius) indicates mean air temperature in chamber, and *P* (mm Hg) indicates instant atmospheric pressure. Cumulative N_2_O emission (E, kg N ha^−1^) was calculated by the equation (Chen et al., 2016),

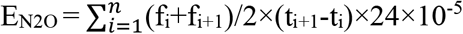

where *f* indicates gas flux (µg N_2_O-N m^-2^ h^−1^), *n* indicates the total measurement numbers, *i* is the sampling time, and *(t*_*i+1*_ *-t*_*i*_ ) represents the interval of two sampling. We measured the N_2_O flux only in the early stage of corn growth because N_2_O emissions usually peaked soon after fertilization (Yang et al., 2017).

### Soil physicochemical analysis

Soil samples were collected on days 0, 15, 60 and 120 after seeding. At each time point, we carefully collected 3 soil cores with a depth of 0-20 cm in each pot about 10 cm away from the axial root and mixed them evenly to make composite samples. In all, 192 soil samples were collected for downstream analysis, and they were stored at 4°C for property measurement and -80°C for DNA extraction, respectively.

Soil pH was measured (water to soil ratio of 2.5:1 v/w) by a Delta 320 pH meter (Mettler-Toledo, China). Soil water content was determined by oven-drying the fresh samples in 105 °C for 24 hours. NH_4_^+^-N and NO_3_^-^-N were extracted with 1 M KCl solution and determined by the continuous flow analysis system (SAN++, Skalar, Holland). A Fusion Total Organic Carbon Analyzer (Tekmar Dohrmann, USA) was used to measure the dissolved organic carbon (DOC) extraction (0.5M K_2_ SO_4_).

Potential nitrification rate (PNR) was determined by the sodium azide inhibition method (Wang et al., 2018a). Fresh soil sample (5g) was added to a 50ml centrifuge tube containing 20ml (1mM) solution of phosphate buffer (PBS) (g L^−1^: NaCl, 8.0; KCl, 0.2; NaHPO_4_, 1.44; KH_2_ PO_4_, 0.24; pH=7.4) with 1mM (NH_4_)_2_ SO_4_ and 10mM KClO_3_ to inhibit nitrite oxidation. After incubating the suspension in dark on a shaker at 180 rpm for 24 hours (25°C), the nitrite was extracted using 5ml of 2M KCl and determined by a Griess-Ilosvay spectrophotometry.

### DNA extraction, Terminal Restriction Fragment Length Polymorphism (T-RFLP) and real-time quantitative PCR (qPCR) analysis

According to the manufacturer’s instructions, soil metagenomic DNA was extracted from 0.3g soil using mobio PowerSOil DNA Isolation Kit (MoBio laboratories, Carlsbad, CA, USA). A Nanodrop ND-2000c UV-Vis spectrophotometer (NanoDrop Technologies, Wilmington, DE, USA) was used to measure the quantity and quality of DNA. The DNA extractions were stored in -20°C freezer for molecular analysis.

T-RFLP was employed to determine the community structure of denitrifying bacteria containing *nirK* and *nosZ* genes. Primer pairs F1aCu/R3Cu and nosZ-F/nosZ1662R were employed for *nirK* gene and *nosZ* gene amplification, respectively (Hallin and Lindgren, 1999; Throback et al., 2004). The forward primers were labelled with fluorescent dye 6-carboxyfluorescein (6-FAM) mark. The 50µl PCR amplification system contains 25µl 2× SYBR Green Premix Ex Taq™ (TaKaRa Biotechnology, Japan), 2µl of each primer (10µM), 3µl of template DNA and 20µl ddH_2_O. PCR cycling conditions of the *nirK* and *nosZ* genes are listed in Table S1. Purified amplicons were digested with restriction enzymes MspI (TaKaRa, Japan) and submitted for fragment scan. Terminal spectra were analyzed by GeneMarker V2.2.0 software (SoftGenetics-LLC, Pennsylvania, USA). The fragments with less than 1bp length variation were combined, and peaks with height less than 0.5% of the maximum value were excluded.

The copies of *amoA, nirK* and *nosZ* genes was determined using an iCycler iQ5 Real-Time PCR detection system (Bio-Rad Laboratories, USA). Primer pairs and thermal cycling programs are listed in Table S1. The 25µl qPCR reaction system contains 12.5 µl 2×SYBR Green Premix Ex Taq™ (TaKaRa Biotechnology, Japan), 0.5µl of each primer (10 µM), 1.5µl template DNA and 10.5µl ddH_2_O.

### Statistical Analysis

One-way analysis of variance (ANOVA) followed by Ducan test (significance level *P* < 0.05) were used to test the difference of gas emissions/soil properties/genes copies among fertilization treatments. The Bray-Curtis dissimilarity matrix was calculated, based on the fragment composition of T-RFLP analysis, to determine the response of denitrifier community containing *nirK* and *nosZ* genes to fertilization treatments. Resultant matrices were ordinated by the nonmetric multidimensional scaling (NMDS) algorithm. These statistical analyses were conducted with SPSS19 software (IBM, USA) and the *vegan* package (Dixon, 2003) in R (www.r-project.org). Boxplots were generated using the *ggplot2* package (Ginestet, 2011) in R. Structural equation model (SEM) was used to evaluate the effects of management, denitrifiers and edaphic properties on the N_2_O flux. Parameters including approximate root mean square error (0≤RMSEA≤0.05), *P* value (> 0.05) and Chi-square were used to evaluate the fitness of the model. The SEM analysis was conducted using AMOS17.0 (AMOS IBM, USA).

## Results

### Soil carbon, nitrogen, pH, and nitrification potential

Compared with the urea fertilization treatment (N), adding DCD (N+NI) significantly (*P*<0.05) increased the NH_4_^+^ in all the soils, and a maximum value was observed around the 15th days after seeding; by contrast, the effect of straw incorporation (N+NI) was weaker (Fig. 1 a-c). Both DCD and straw incorporation decreased soil NO_3_^-^, and the former had stronger influence (Fig. 1d-f). Compared with N treatment, DCD or straw didn’t significantly affect dissolved organic carbon (DOC) in all soils (Fig. S2). Fertilizations (N, N+NI, N+SI) caused soil acidification and after 60 days, the lowest pH appeared in the red (4.93) and black (4.35) soils (Fig S3). The potential nitrification rate (PNR) first decreased constantly after 15 days in the fluvo-aquic soil but may retrieve in the other two (Fig S4). DCD significantly decrease PNR only in the black soil, and straw incorporation had no consistent effect.

**Fig. 1.**
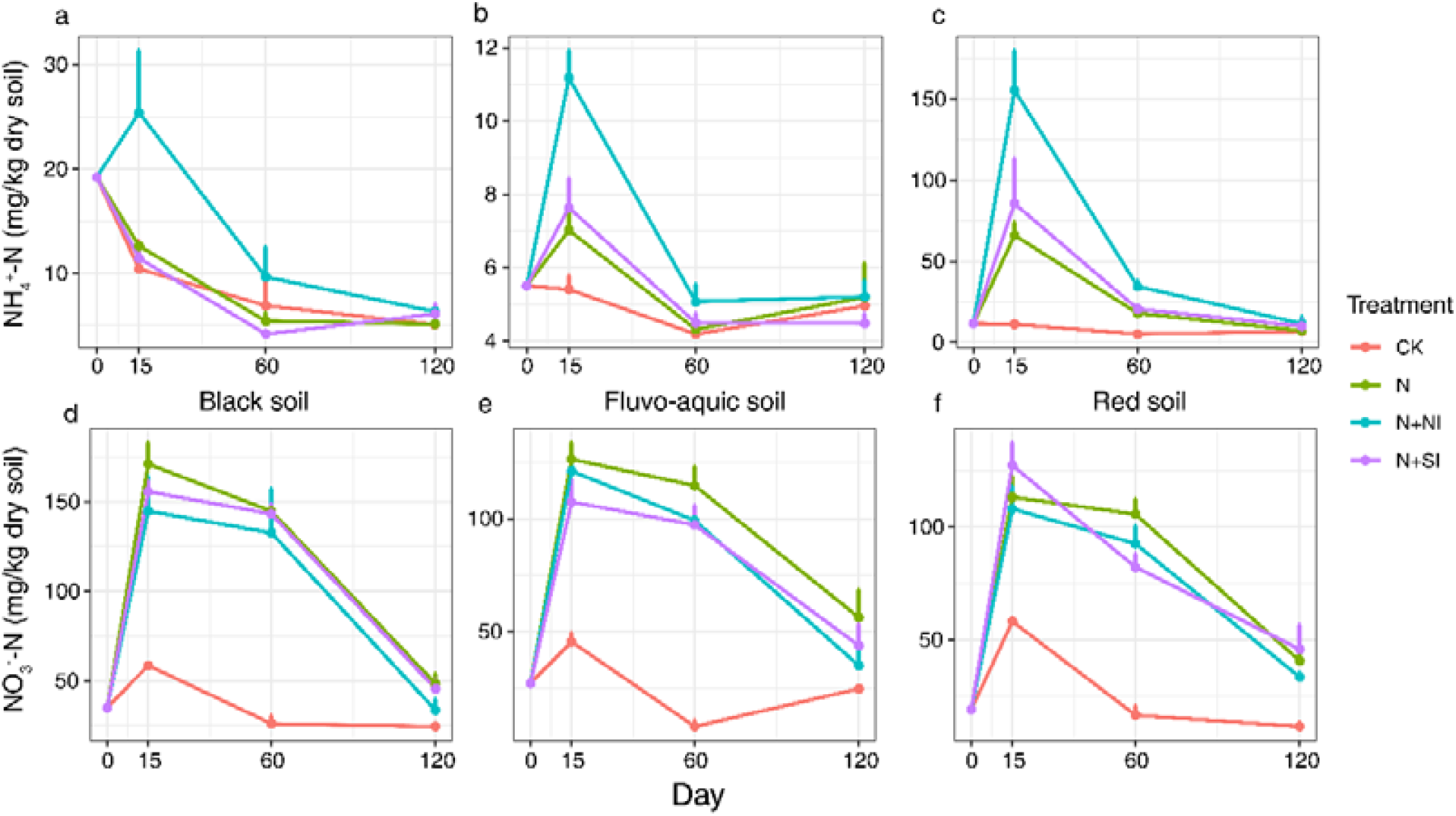
Soil ammonia and nitrate dynamics in the black, fluvo-aquic and red soils under different fertilization treatments (**a**-**f**). Error bars present standard deviations of means (n = 4).

### Flux and accumulative N_2_O emissions

In the black soil, the fertilized treatments (N, N+NI, N+SI) had higher N_2_O flux than the unfertilized control (CK) (Fig S1a) and therefore accumulated more N_2_O (Fig 2a). Straw incorporation increased ∼50% N_2_O emissions from black soil, while DCD had no significant effect. In the fluvo-aquic soil, nitrogen fertilization (N) significantly increased the N_2_O flux at the early stage (days 0-3); adding DCD rather than straw incorporation can effectively alleviate such trend (Fig S1b). Accumulatively, DCD reduced ∼60% N_2_O emission from the fluvo-aquic soil, while straw incorporation had no such strong effect (Fig 2b). DCD had no significant effect on the N_2_O flux (Fig S1.c) or accumulative emissions (Fig 2c) in the red soil, while straw incorporation strongly and significantly increased the N_2_O flux throughout the whole stage, and such effect became stronger with time (*P*<0.05). Specifically, the N_2_O accumulation of N+SI treatment was ∼600% higher than N treatments (*P*<0.05).

**Fig. 2.**
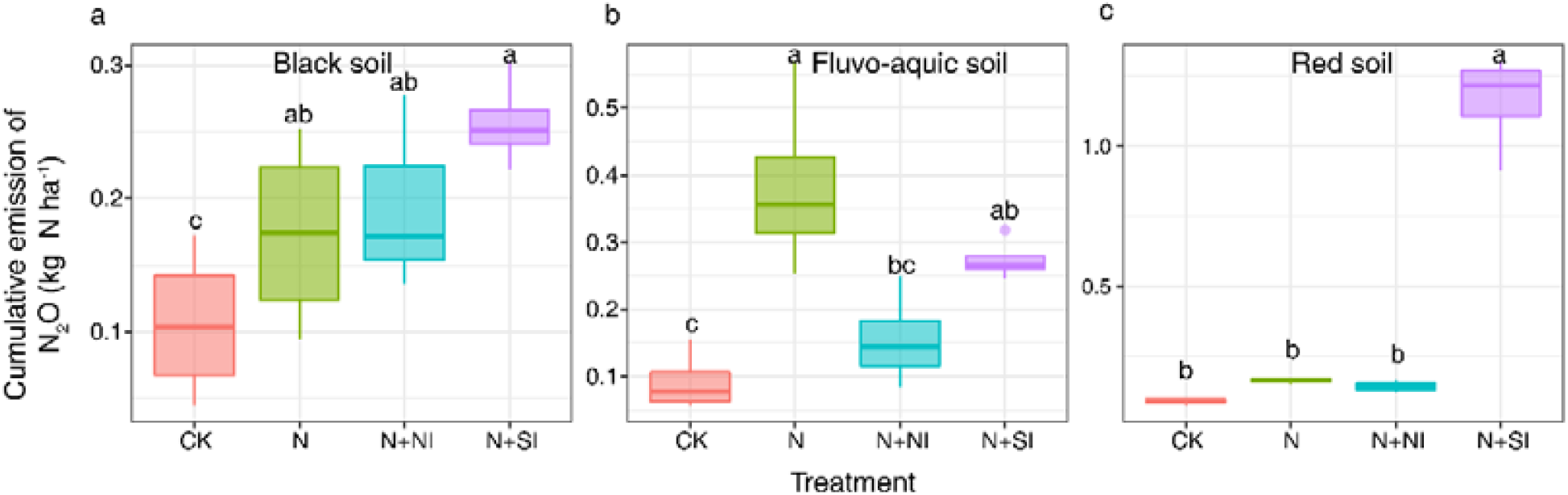
The accumulative N_2_O emission during the early growth stage of corn under different fertilization treatments (**a**-**c**). Error bars present standard deviations of means (n = 4). The different lowercase letters indicate significant difference among treatments by Duncan’s multiple range test (*P* < 0.05).

### *nirK*- and *nosZ*-containing denitrifier community

T-RFLP analysis showed that the black soil, fluvo-aquic soil and red soil had distinct denitrifier communities (Fig. 3). Both the *nirK*-containing denitrifiers assemblages and *nosZ*-containing denitrifiers were similar within the same soil but distinct between the three soils. And *nosZ*-containing denitrifier community is more conservative than *nirK*-containing denitrifier within the same soil. No significant difference was observed among different treatments on the same soil; either DCD or straw incorporation did not change the denitrifier community.

**Fig. 3.**
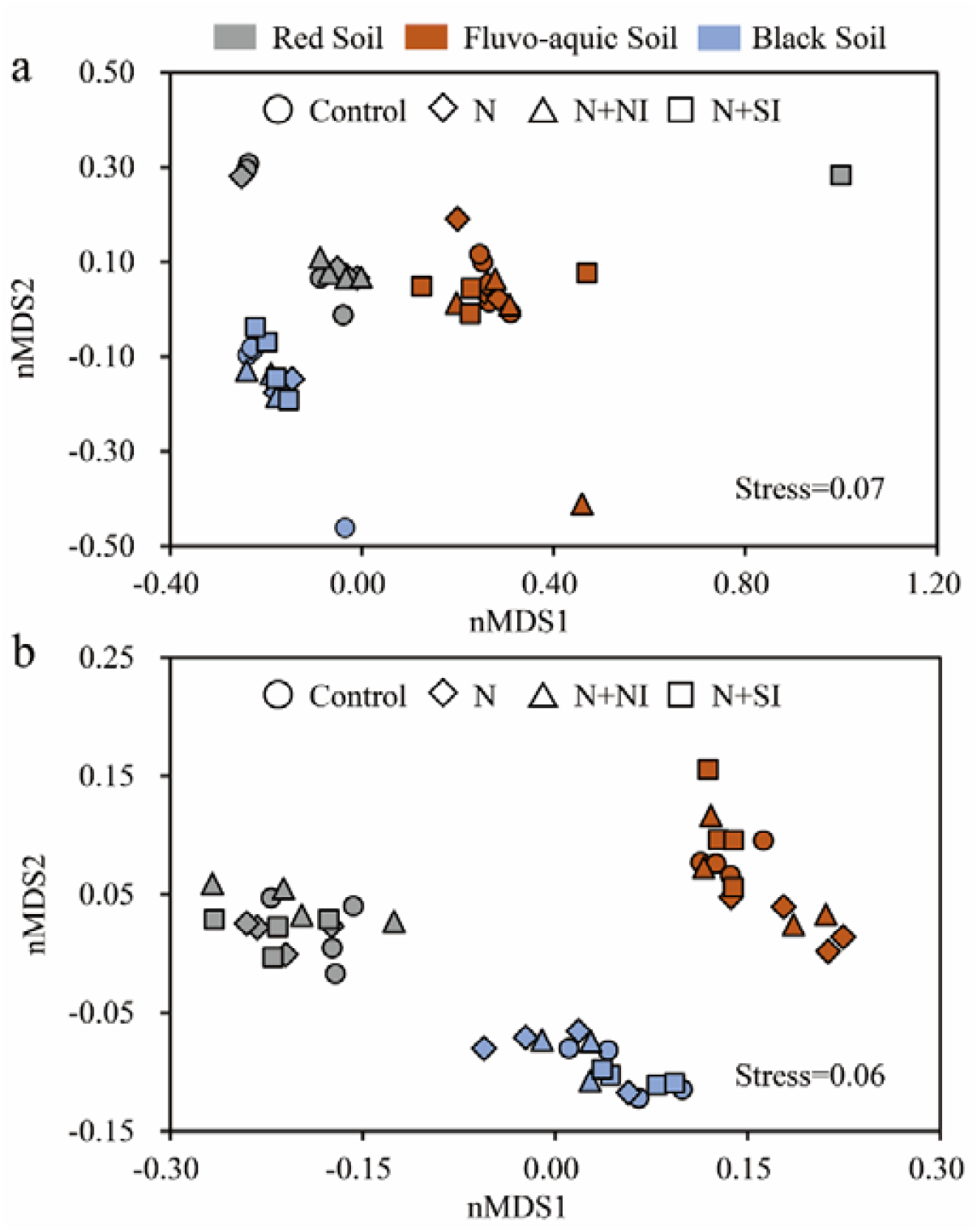
Non-metric multidimensional scaling (nMDS) of *nirK* (**a**) and *nosZ* (**b**) gene-containing denitrifiers based on the Bray-Curtis dissimilarity matrix of T-RFs among three soil types

### Abundances of nitrifier and denitrifier functional genes

The abundance of archaeal and bacterial *amoA* gene (copies *per* gram dry soil) were both lower in the red soil (4.56 × 10^6^ to 6.29 × 10^6^ for AOA, and 1.79 × 10^7^ to 2.53 × 10^7^ for AOB) than in the fluvo-aquic soil (3.76 × 10^7^ to 1.13 × 10^8^ for AOA, and 4.23 × 10^7^ to 5.75 × 10^7^ for AOB), but no significant changes of the *amoA* genes copies was observed among different fertilization treatments (Fig. S5).

Overall, the abundance of *nirK* gene in the black soil (4.3 × 10^6^ to 7.57 × 10^6^) was lower than in the fluvo-aquic soil (2.98 × 10^7^ to 4.37 × 10^7^) and the red soil (3.94 × 10^7^ to 6.22 × 10^7^); however, for each soil, there was no significant changes in *nirK* gene abundance among different fertilization treatments (Fig. 4a-c). DCD and straw incorporation increased the abundance of *nosZ* gene in the black and fluvo-aquic soils (Fig. 4d). No significant variation of the *nosZ* gene abundances was observed among all treatments in the red soil (Fig. 4 e-f).

**Fig. 4.**
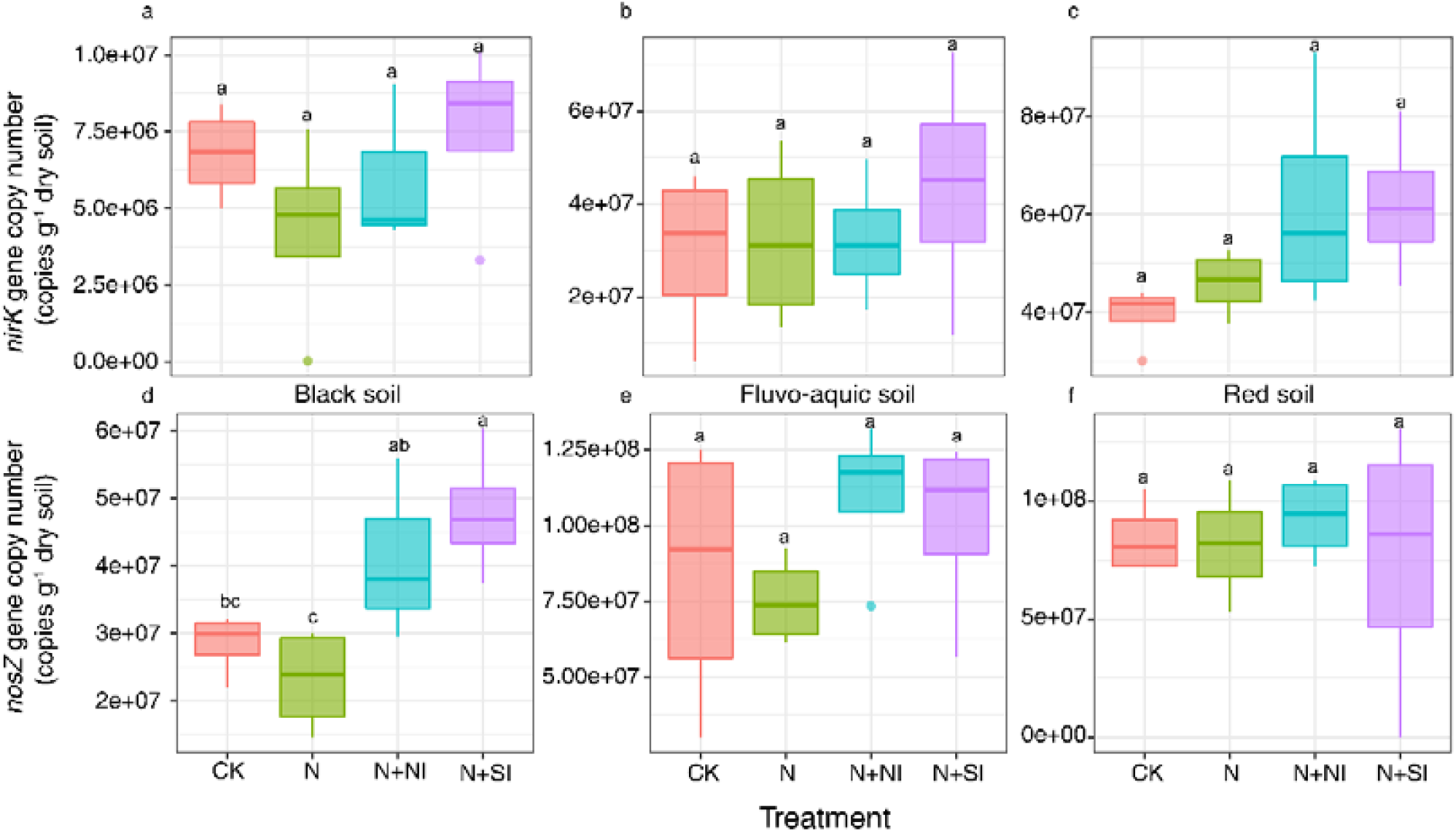
The abundance of denitrifiers (based on the copies of *nirK* and *nosZ* genes) in black, fluvo-aquic and red soils under different fertilization treatments (**a**-**f**). Error bars present standard deviations of means (n = 4). The different lowercase letters indicate significant difference among treatments Duncan’s multiple range test (*P* < 0.05)

### Structural equation modelling

To further clarify their contributions to N_2_O emission across different soils, we used the structural equation model (SEM) to evaluate the effects (both direct and indirect) of straw incorporation, DCD, soil dissolved organic carbon, nitrite reductase and nitrous oxide reductase. Dissolved organic carbon (0.72) and straw incorporation (0.47) had strong and positive effects on N_2_O flux, whereas nitrous oxide reductase gene (*nosZ*) abundance had significantly negative effect (−0.52) on N_2_O flux (Fig. 5a). Straw incorporation, however, does not directly increase the dissolved organic carbon, which is an important factor that regulates the *nirK* gene abundance and communities containing *nosZ* gene. Accounting all the biotic and abiotic factors together, the SEM explained 51.0% of the variance in N_2_O flux (Fig. 5a). It supported that straw incorporation is an important management practice that can increase the risk of N_2_O emission; soil DOC is the most important abiotic factor while *nosZ* gene abundance is the only significant biotic factor that affect the N_2_O emission (Fig. 5b).

**Fig. 5.**
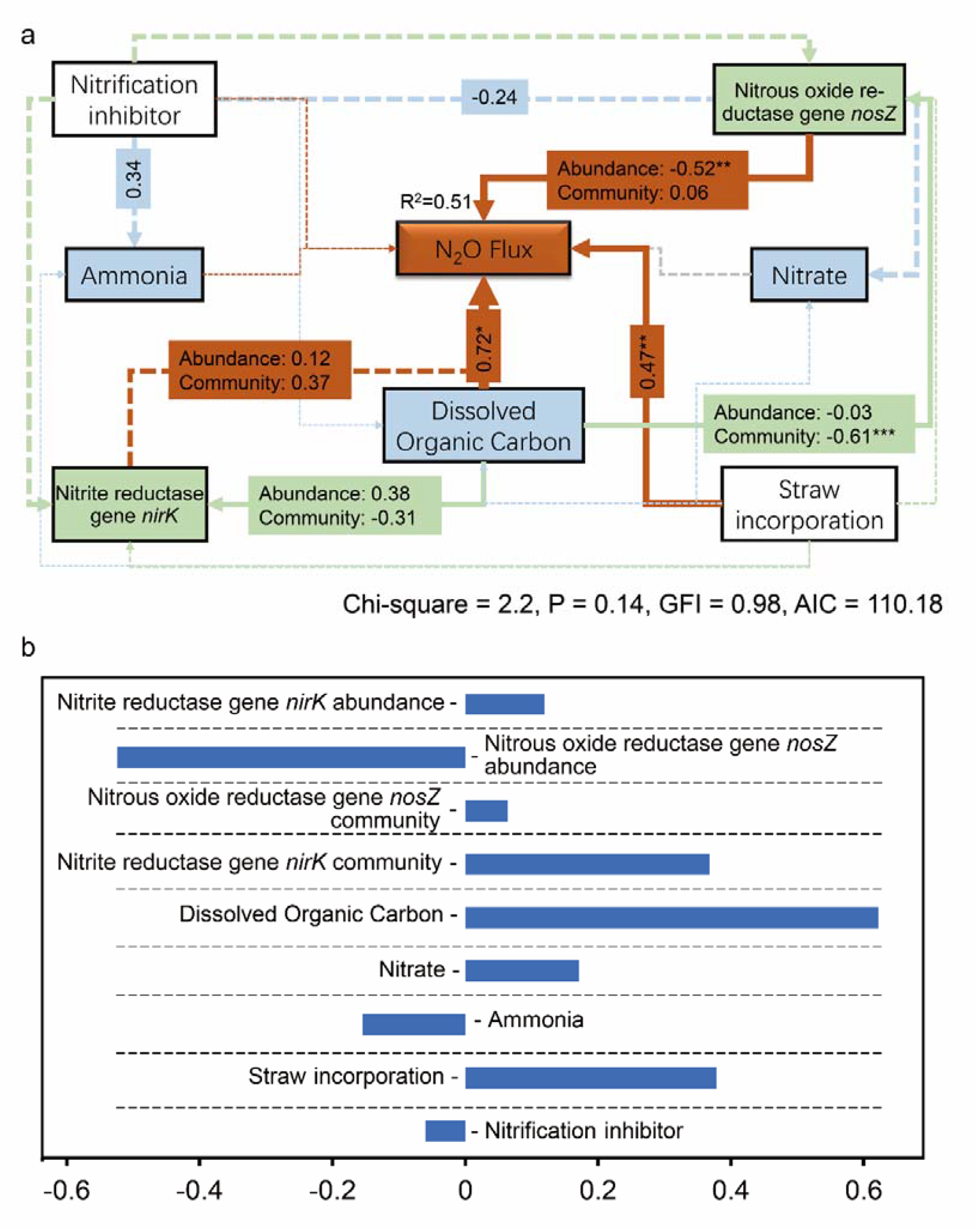
(**a**) The structural equation model exhibiting the effects of abiotic and biotic factors on N_2_O flux. Numbers adjacent to arrows are indicative of the effect-size of the relationship. * indicates *P* < 0.05; ** indicates *P* < 0.01; *** indicates *P* < 0.001. Continuous and dashed lines indicate significant and non-significant relationships, respectively. Orange, blue and grey lines indicate the colors of variables the arrows pointed to, respectively. The width of arrows is proportional to the strength of path coefficients. R^2^ denotes the proportion of variance explained by the model. (**b**) Standardized total effects (direct plus indirect effects) derived from the structural equation models depicted above.

## Discussion

In facing of the increasing concentration of atmospheric N_2_O, various management practices were used to improve nitrogen fertilizer utilization and regulate N_2_O emissions from cropland soils (Zhu et al. 2019). Nitrification inhibitors like DCD can inhibit the conversion from NH_4_^+^-N to NO_2_^-^/NO_3_^-^and thereafter reduce substrate from pathways generating N_2_O. Straw incorporation can significantly increase soil organic carbon (SOC) by 12.8% and improve soil nutrient availability for crop in global croplands (Liu et al., 2014), and can potentially mitigate N_2_O emissions (Badagliacca et al., 2017). However, the performance of DCD and straw incorporation on N_2_O emission from different types of soils had not been parallelly evaluated. Using controlled experiment and DNA-based molecular analysis on three distinct cropland soils, we showed that the performance of DCD and straw incorporation on N_2_O emission are distinct in different soils, and multiple factors contributed to such pattern.

### Soil physicochemical properties and N_2_O emission

Soil pH is reported to be important factor affecting N_2_O emissions (Signor and Cerri, 2013), and some studies showed that pH can directly affects the activity of nitrification and denitrification reductase (Granli and Bockman, 1994). A few studies reported that the emission of N_2_O would be greater in lower pH habitat (Chapuis-Lardy et al., 2007). Our study supported this since straw incorporation increased more N_2_O emission in low pH red soils (pH=4.43) than high pH fluvo-aquic soils (pH=7.18).

Besides soil pH, the organic carbon is another important regulating factor. Soil carbon not only directly provides energy to the denitrifier, but also promotes microbial consumption on O_2_, creating favorable conditions for denitrification (Beauchamp E G, 1989). Miller et al. (2008) found that the available carbon increases microbial activity and oxygen consumption, leading to favorable conditions for denitrification. Previous studies have found that increased DOC levels are associated with increased N_2_O emissions, as activated carbon is a key role for the electron acceptor for microbial denitrification (Harter et al., 2014). Besides, some studies reported that DOC is a measurement of existing resources for microbial growth and biodegradation and is generally considered a good indicator of carbon effectiveness (Jensen et al., 1997; Liang et al., 1996). Many others reported that short-term straw incorporation to the field increases the DOC concentration than not incorporation it to the field (Chen et al., 2017; Guo et al., 2014; Li et al., 2013). Our study supported that straw incorporation significantly promoted DOC content in all soils. The decomposition of straw releases nutrients and DOC, which stimulates the biological activities of microorganisms (Wang et al., 2015).

### Response of functional microbes

The *nirK* gene encodes nitrite reductase that catalyzes the conversion from NO_2_^-^ to NO, while the *nosZ* gene encodes nitrous oxide reductase that can consume N_2_O (N_2_O-N_2_), the only known biological link that reduces N_2_O emissions. DCD affected the abundance of the *nirK* genes, probably because the *nirK* gene communities are also ammonia oxidizers, which are inhibited by DCD (Di et al., 2014). Huang et al. (2004) reported that denitrifying bacteria may consume soil DOC, thereby producing denitrification and increasing N_2_O emissions. Our research found that straw incorporation significantly increased N_2_O emissions especially in red soils (Fig. 2). The nitrogen in the straw was returned to the field and became an additional nitrogen input, which might be the main cause of the direct increase in N_2_O emissions (Lu et al., 2010).

Nitrification inhibitors is a widely used strategy to reduce soil N_2_O emissions (Snyder et al., 2009) since it decreased the autotrophic nitrification (Bauhus et al., 1996). Some studies reported that DCD had a significant effect on reducing N_2_O emission of maize compared with only nitrogen application (Ding et al., 2011; Zhou et al., 2016). Our result is consistent with the above findings and showed that the N+NI treatment reduced N_2_O emission in fluvo-aquic soil compared with the N treatment (Fig. 2b). Compared with other treatment, N+NI treatment increased the concentration of NH_4_^+^-N (Fig. S1), possibly because nitrification inhibitor application blocked ammonia monooxygenase enzyme and retained NH_4_^+^-N (Abbasi and Adams, 2000; Wolt, 2004), thereby mitigating N_2_O emissions from soils.

## Conclusions

Our study provided evidence that straw incorporation better suits the fluvo-aquic soil but might not be suitable for the black or red soils since it can potentially increase N_2_O emission (especially from the red soil). Nitrification inhibitor DCD is comparably safe in N_2_O emissions and could be deployed in variable soils. The response of *nosZ* gene abundance to inhibitor/straw incorporation is the main biological factor determining the soil N_2_O emission from different soils. This work advanced our understanding on the agricultural practices and N_2_O emission in cropland soils, suggesting that management practices should be used as *per* soil types to balance between nitrogen use efficiency and N_2_O emission.

## Acknowledgements

This work was financially supported by the National Key R&D Program (2017YFD0800604) and National Natural Science Foundation of China (41771288). We thank Peng-Xia Xu, Ya-Qi Wang, Shuai Du, and Liu-Ying Mo for their assistance in greenhouse and laboratory work.

## Author contributions

LMZ and JZH designed the study. FL collected data in field and lab work. YBZ and JTW analyzed the data. YBZ, JTW, HWH and LMZ wrote the manuscript in close consultation from all authors.

## References

Abbasi, M.K., Adams, W.A., 2000. Estimation of simultaneous nitrification and denitrification in grassland soil associated with urea-N using 15N and nitrification inhibitor. Biology and Fertility of Soils. 31, 38–44.

Badagliacca, G., Ruisi, P., Rees, R.M., Saia, S., 2017. An assessment of factors controlling N_2_O and CO_2_ emissions from crop residues using different measurement approaches. Biology and Fertility of Soils. 53, 547–561.

Bauhus, J., Meyer, A.C., Brumme, R., 1996. Effect of the inhibitors nitrapyrin and sodium chlorate on nitrification and N_2_O formation in an acid forest soil. Biology and Fertility of Soils. 22, 318–325.

Beauchamp E G, Trevors J T, Paul J W, 1989. Carbon sources for bacterial denitrification. Advances in Soil Science. 10, 113–142.

Butterbach-Bahl, K., Baggs, E.M., Dannenmann, M., Kiese, R., Zechmeister-Boltenstern, S., 2013. Nitrous oxide emissions from soils: how well do we understand the processes and their controls? Philosophical Transactions of The Royal Society B Biological Sciences. 368, 20130122.

Chapuis-Lardy, L., Wrage, N., Metay, A., Chotte, J.-L., Bernoux, M., 2007. Soils, a sink for N_2_O? A review. Global Change Biology. 13, 1–17.

Chen, H., X. Li, F. Hu, and W. Shi. 2013. Soil nitrous oxide emissions following crop residue addition: a meta-analysis. Global Change Biology. 19, 2956–2964.

Chen, Z., Wang, H., Liu, X., Zhao, X., Lu, D., Zhou, J., Li, C., 2017. Changes in soil microbial community and organic carbon fractions under short-term straw return in a rice–wheat cropping system. Soil and Tillage Research. 165, 121–127.

Di, H.J., Cameron, K.C., Podolyan, A., Robinson, A., 2014. Effect of soil moisture status and a nitrification inhibitor, dicyandiamide, on ammonia oxidizer and denitrifier growth and nitrous oxide emissions in a grassland soil. Soil Biology & Biochemistry. 73, 59–68.

Di, H.J., Cameron, K.C., Sherlock, R.R., 2007. Comparison of the effectiveness of a nitrification inhibitor, dicyandiamide, in reducing nitrous oxide emissions in four different soils under different climatic and management conditions. Soil Use and Management. 23, 1–9.

Ding, W.X., Chen, Z.M., Yu, H.Y., Luo, J.F., Yoo, G.Y., Xiang, J., Zhang, H.J., Yuan, J.J., 2015. Nitrous oxide emission and nitrogen use efficiency in response to nitrophosphate, N-(n-butyl) thiophosphoric triamide and dicyandiamide of a wheat cultivated soil under sub-humid monsoon conditions. Biogeosciences. 12, 803–815.

Ding, W.X., Yu, H.Y., Cai, Z.C., 2011. Impact of urease and nitrification inhibitors on nitrous oxide emissions from fluvo-aquic soil in the North China Plain. Biology and Fertility of Soils. 47, 91–99.

Dixon, P., 2003. VEGAN, a package of R functions for community ecology. Journal of Vegetation Science. 14, 927–930.

Ginestet, C., 2011. ggplot2: Elegant Graphics for Data Analysis. Journal of the Royal Statistical Society Series a-Statistics in Society. 174, 245–245.

Granli, T., Bockman, O.C., 1994. Nitrous oxide (N_2_O) from agriculture. Third Congress of the European Society for Agronomy, Proceedings, 800–801.

Guo, L.J., Zhang, Z.S., Wang, D.D., Li, C.F., Cao, C.G., 2014. Effects of short-term conservation management practices on soil organic carbon fractions and microbial community composition under a rice-wheat rotation system. Biology and Fertility of Soils. 51, 65–75.

Hallin, S., Lindgren, P.E., 1999. PCR detection of genes encoding nitrile reductase in denitrifying bacteria. Applied and Environmental Microbiology. 65, 1652–1657.

Harter, J., Krause, H.M., Schuettler, S., Ruser, R., Fromme, M., Scholten, T., Kappler, A., Behrens, S., 2014. Linking N_2_O emissions from biochar-amended soil to the structure and function of the N-cycling microbial community. The ISME journal. 8, 660–674.

Hu, H.W., Chen, D., He, J.Z., 2015. Microbial regulation of terrestrial nitrous oxide formation: understanding the biological pathways for prediction of emission rates. FEMS microbiology reviews . 39, 729–749.

Hu, H.W., Zhang, L.M., Yuan, C.L., Zheng, Y., Wang, J.T., Chen, D., He, J.Z., 2015. The large-scale distribution of ammonia oxidizers in paddy soils is driven by soil pH, geographic distance, and climatic factors. Frontiers in Microbiology. 6:938.

Huang, Y., Zou, J., Zheng, X., Wang, Y., Xu, X., 2004. Nitrous oxide emissions as influenced by amendment of plant residues with different C:N ratios. Soil Biology & Biochemistry. 36, 973–981.

IPCC, 2013. Climate Change 2013: The Physical Science Basis. Contribution of Working Group I to the Fifth Assessment Report of the Intergovernmental Panel on Climate Change [Stocker, T.F., D. Qin, G.-K. Plattner, M. Tignor, S.K. Allen, J. Boschung, A. Nauels, Y. Xia, V. Bex and P.M. Midgley (eds.)]. Cambridge University Press, Cambridge, United Kingdom and New York, NY, USA 1535PP.

IPCC, 2014. Climate Change 2014: Synthesis Report. Contribution of Working Groups I, II and III to the Fifth Assessment Report of the Intergovernmental Panel on Climate Change.[Core Writing Team, R.K. Pachauri and L.A. Meyer (eds.)]. IPCC.Geneva, Switzerland 151 pp.

Jensen, L.S., Mueller, T., Magid, J., Nielsen, N.E., 1997. Temporal variation of C and N mineralization, microbial biomass and extractable organic pools in soil after oilseed rape straw incorporation in the field. Soil Biology & Biochemistry. 29(7), 1043–1055.

Jones, C. M., and S. Hallin. 2010. Ecological and evolutionary factors underlying global and local assembly of denitrifier communities. The ISME Journal. 4, 633–641.

Lam, S.K., Suter, H., Mosier, A.R., Chen, D., 2017. Using nitrification inhibitors to mitigate agricultural N_2_O emission: a double-edged sword? Global Change Biology. 23, 485–489.

Li, F., Cao, X., Zhao, L., Yang, F., Wang, J., Wang, S., 2013. Short-term effects of raw rice straw and its derived biochar on greenhouse gas emission in five typical soils in China. Soil Science and Plant Nutrition. 59, 800–811.

Liang, B.C., Gregorich, E.G., Schnitzer, M., Voroney, R.P., 1996. Carbon mineralization in soils of different textures as affected water-soluble organic carbon extracted from composted dairy manure. Biology and Fertility of Soils. 21, 10–16.

Liu, C., Lu, M., Cui, J., Li, B., Fang, C., 2014. Effects of straw carbon input on carbon dynamics in agricultural soils: a meta-analysis. Global Change Biology. 20, 1366–1381.

Lu, F., Wang, X., Han, B., Ouyang, Z., Duan, X., Zheng, H., 2010. Net mitigation potential of straw return to Chinese cropland: estimation with a full greenhouse gas budget model. Ecological Applications. 20, 634–647.

Meng, X., Li, Y., Yao, H., Wang, J., Dai, F., Wu, Y., Chapman, S., 2020. Nitrification and urease inhibitors improve rice nitrogen uptake and prevent denitrification in alkaline paddy soil. Applied Soil Ecology. 154.

Miller, M. N., C. E. Dandie, B. J. Zebarth, D. L. Burton, C. Goyer, and J. T. Trevors. 2012. Influence of carbon amendments on soil denitrifier abundance in soil microcosms. Geoderma. 170:48–55.

Miller, M.N., Zebarth, B.J., Dandie, C.E., Burton, D.L., Goyer, C., Trevors, J.T., 2008. Crop residue influence on denitrification, N_2_O emissions and denitrifier community abundance in soil. Soil Biology and Biochemistry. 40, 2553–2562.

Ning, J., Ai, S., Cui, L., 2018. Dicyandiamide has more inhibitory activities on nitrification than thiosulfate. PLoS One. 13, e0200598.

Ravishankara, A.R., Daniel, J.S., Portmann, R.W., 2009. Nitrous Oxide (N_2_O): The Dominant Ozone-Depleting Substance Emitted in the 21st Century. Science. 326, 123–125.

Signor, D., Cerri, C.E.P., 2013. Nitrous oxide emissions in agricultural soils: a review. Pesquisa Agropecuária Tropical. 43, 322–338.

Snyder, C.S., Bruulsema, T.W., Jensen, T.L., Fixen, P.E., 2009. Review of greenhouse gas emissions from crop production systems and fertilizer management effects. Agriculture Ecosystems & Environment 133, 247–266.

Throback, I.N., Enwall, K., Jarvis, A., Hallin, S., 2004. Reassessing PCR primers targeting nirS, nirK and nosZ genes for community surveys of denitrifying bacteria with DGGE. Fems Microbiology Ecology. 49, 401–417.

Wang, H., Liu, S., Zhang, X., Mao, Q., Li, X., You, Y., Wang, J., Zheng, M., Zhang, W., Lu, X., Mo, J., 2018a. Nitrogen addition reduces soil bacterial richness, while phosphorus addition alters community composition in an old-growth N-rich tropical forest in southern China. Soil Biology and Biochemistry. 127, 22–30.

Wang, W., Lai, D.Y.F., Wang, C., Pan, T., Zeng, C., 2015. Effects of rice straw incorporation on active soil organic carbon pools in a subtropical paddy field. Soil and Tillage Research. 152, 8–16.

Wang, Y.Q., Bai, R., Di, H.J., Mo, L.Y., Han, B., Zhang, L.M., He, J.Z., 2018b. Differentiated Mechanisms of Biochar Mitigating Straw-Induced Greenhouse Gas Emissions in Two Contrasting Paddy Soils. Frontiers in Microbiology. 9, 2566.

Wolt, J.D., 2004. A meta-evaluation of nitrapyrin agronomic and environmental effectiveness with emphasis on corn production in the Midwestern USA. Nutrient Cycling in Agroecosystems. 69, 23–41.

Wu, L., Hu, R., Tang, S., Shaaban, M., Zhang, W., Shen, H., Xu, M., 2020. Nitrous oxide emissions in response to straw incorporation is regulated by historical fertilization. Environmental Pollution. 266, 115292.

Xu, C., Han, X., Ru, S., Cardenas, L., Rees, R.M., Wu, D., Wu, W., Meng, F., 2019. Crop straw incorporation interacts with N fertilizer on N_2_O emissions in an intensively cropped farmland. Geoderma. 341, 129–137.

Yang, L., Zhang, X., Ju, X., 2017. Linkage between N_2_O emission and functional gene abundance in an intensively managed calcareous fluvo-aquic soil. Scientific Reports. 7, 43283.

Zhao, X., B.-Y. Liu, S.-L. Liu, J.-Y. Qi, X. Wang, C. Pu, S.S. Li, X.Z. Zhang, X.G. Yang, R. Lal, F. Chen, and H.-L. Zhang. 2020. Sustaining crop production in China’s cropland by crop residue retention: A meta-analysis. Land Degradation & Development. 31, 694–709.

Zhou, X., S. Wang, S. Ma, X. Zheng, Z. Wang, and C. Lu. 2020. Effects of commonly used nitrification inhibitors-dicyandiamide (DCD), 3, 4-dimethylpyrazole phosphate (DMPP), and nitrapyrin-on soil nitrogen dynamics and nitrifiers in three typical paddy soils. Geoderma. 380.

Zhou, Y., Zhang, Y., Tian, D., Mu, Y., 2016. Impact of dicyandiamide on emissions of nitrous oxide, nitric oxide and ammonia from agricultural field in the North China Plain. Journal of Environmental Sciences-China. 40, 20–27.

Zhu, G. D., X. T. Song, X. T. Ju, J. B. Zhang, C. Muller, R. Sylvester-Bradley, R. E. Thorman, I. Bingham, and R. M. Rees. 2019. Gross N transformation rates and related N_2_O emissions in Chinese and UK agricultural soils. Science of the Total Environment.∼ 666, 176–186.

