## Supplementary for "Effect of straw incorporation and nitrification inhibitor on nitrous oxide emission in various cropland soils and its microbial mechanism"

**Table S1** Primers and PCR reaction conditions for T-RFLP and qPCR analysis

| Target group | Primers for qPCR | Sequence (5´-3´) | Amplification condition | Reference |
| --- | --- | --- | --- | --- |
| Archaeal *amoA* | Arch-amoAF | STAATGGTCTGGCTTCTTC | 95℃ for 3min; 35 circles of (95℃ for 10s, 55℃ for 30s,72℃for 60s and plate read at 83℃ for 10s) | (Francis et al. 2005) |
|  | Arch-amoAR | GCGGCATCCATCTGTATGT |  |  |
| Bacterial amoA | amoA-1F | GGGGTTTCTACTGG TGG | 95℃ for 3min; 35 circles of (95℃ for10s, 55℃ for 30s,72℃for 60s and plate read at 83℃ for 10s) | (Rotthauwe et al. 1997) |
|  | amoA-2R | TACAACTGGGTGAA CTA |  |  |

|  | FlaCu | ATCATGGTSCTGCCGCG | 95℃ for 3min (1cycle); 95℃ for 30s, 63℃ for 30s(-1℃/cycle), 72℃ for 30s (6 cycles); 95℃ for 30s, 58℃ for 30s, 72℃ for 30s, plate read at 83℃ for 10s (32 cycles)，72℃ 10min | (Hallin and Lindgren 1999) |
| --- | --- | --- | --- | --- |
| *nirK* |  |  |  |  |
|  | R3Cu | GCCTCGATCAGRTTGTGGTT |  |  |
| *nosZ* | nosZ-F | CGYTGTTCMTCGACAGCCAG | 94℃ for 2min; 94℃ for 30s, 57℃ for 30s, (-1℃/cycle), 72℃ for 45s (6 cycles); 94℃ for 30s, 52℃ for 30s, 72℃ for 45s (30 cycles), 72℃ 10min | (Throback et al. 2004) |
|  | NosZ1662R | CGSACCTTSTTGCCSTYGCG |  |  |

| Target group | Primers for qPCR | Sequence (5´-3´) | Thermal profile | Reference |
| --- | --- | --- | --- | --- |
|  | FlaCu | ATCATGGTSCTGCCGCG | 95℃ for 3min (1cycle); 95℃ for 30s, 63℃ for 30s(-1℃/cycle), 72℃ for 30s (6 cycles); 95℃ for 30s, 58℃ for 30s, 72℃ for 30s, plate read at 83℃ for 10s (32 cycles) | (Hallin and Lindgren 1999) |
| *nirK* |  |  |  |  |
|  | R3Cu | GCCTCGATCAGRTTGTGGTT |  |  |
|  | nosZ2F | CGCRACGGCAASAAGGTSMSSGT | 95℃ for 15min (1cycle); 95℃ for 15s, 65℃ for 30s, (-1℃/cycle), 72℃ for 30s (6 cycles); 95℃ for 15s, 60℃ for 15s, 72℃ for 30s, plate read at 80℃ for 15s (40 cycles) | (Henry et al. 2006) |
| *nosZ* |  |  |  |  |
|  | nosZ2R | CAKRTGCAKSGCRTGGCAGAA |  |  |

**Fig. S1** Soil ammonia and nitrate dynamics in the black, fluvo-aquic and red soils under different fertilization treatments (**a**-**f**). Error bars present standard deviations of means (n = 4). The different lowercase letters indicate significant difference among treatments by Duncan's multiple range test (*P* < 0.05)

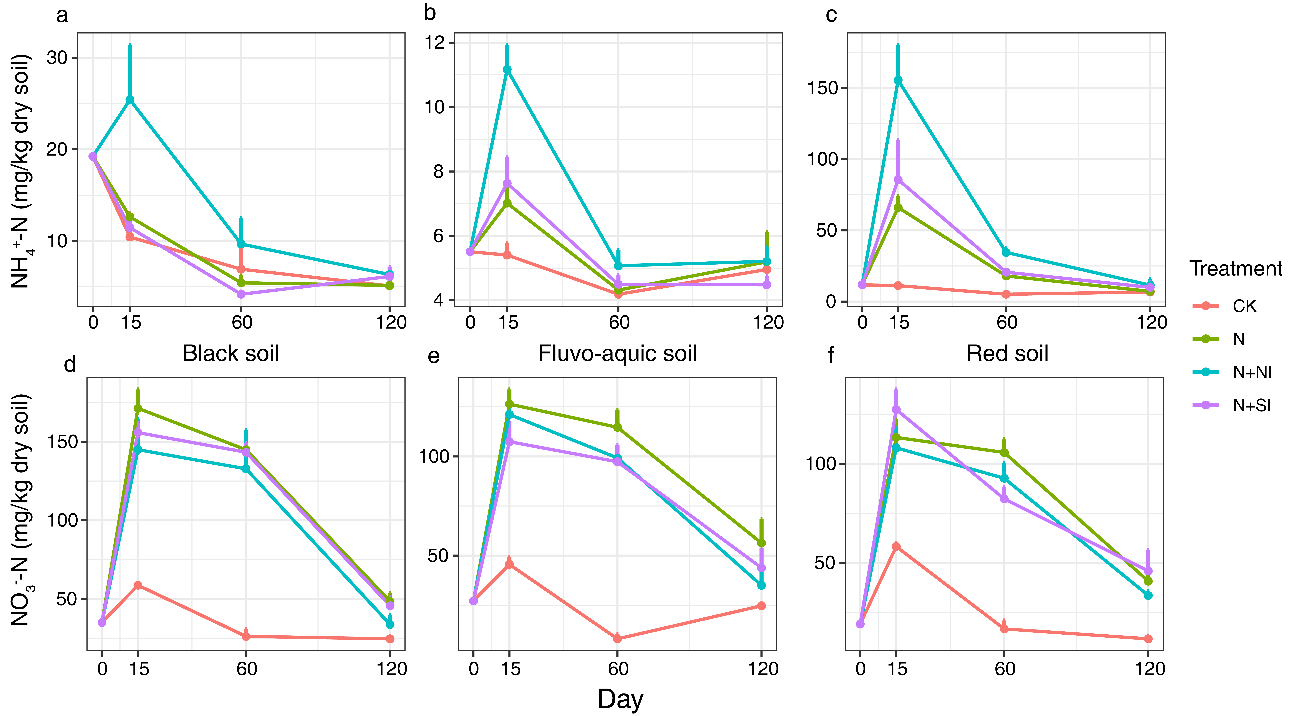

**Fig. S2** Soil dissolved organic carbon (DOC) in black, fluvo-aquic and red soils across different treatments are exhibited in (**a**-**c**). Only day 15 data were analyzed during the whole corn growing season. Error bars present standard deviations of means (n = 4). The different lowercase letters indicate significant difference among different treatments, which was analyzed by Duncan's multiple range test (*P* < 0.05)

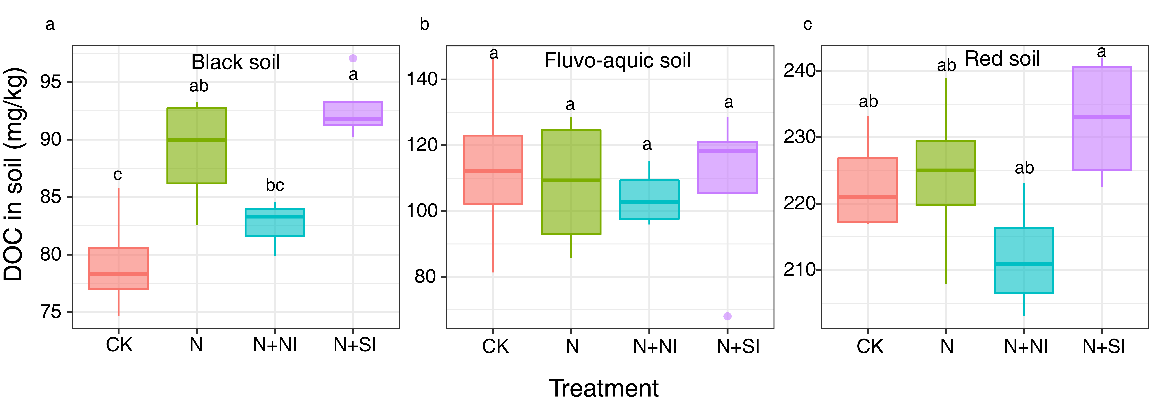

Fig. S3 Soil pH in black, fluvo-aquic and red soils under different fertilization treatments (a-c). Error bars present standard deviations of means (n = 4). The different lowercase letters indicate significant difference among four treatments by Duncan's multiple range test (P < 0.05).

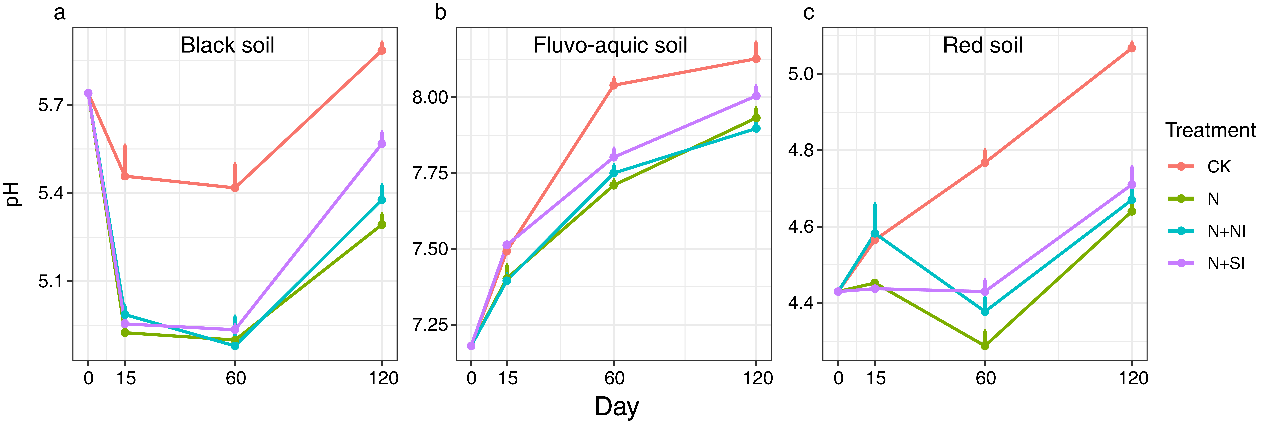

Fig. S4 Potential nitrification rates (PNR) in the black, fluvo-aquic and red soils under different fertilization treatments (a-c). Error bars present standard deviations of means (n = 4). The different lowercase letters indicate significant difference among four treatments by Duncan's multiple range test (P < 0.05).

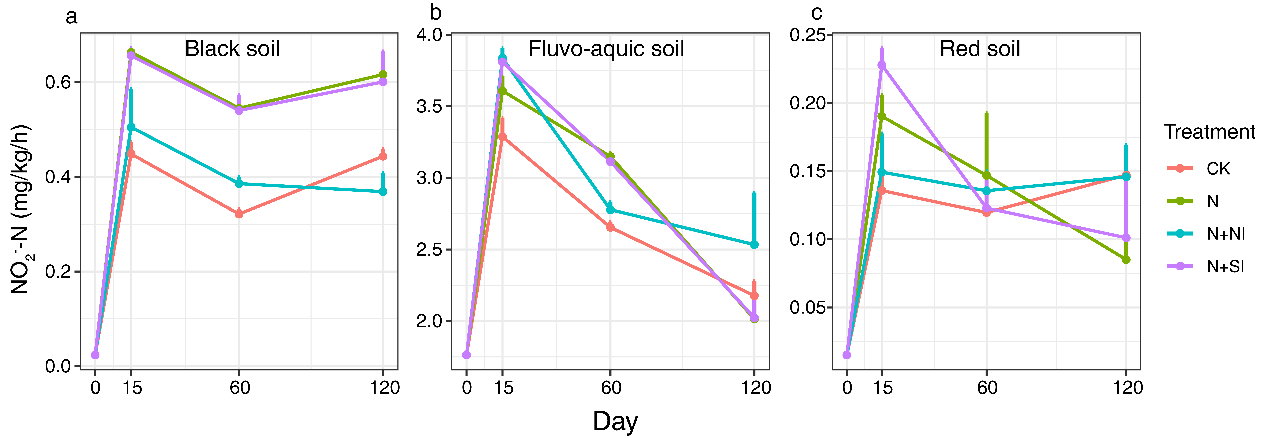

**Fig. S5** The abundance of nitrifiers (based on the *amoA* genes copies of archaea AOA and bacteria AOB) at day 15 after sowing in the fluvo-aquic and the red soils under different fertilization treatments (**a**-**d**). Error bars present standard deviations of means (n = 4). The different lowercase letters indicate significant difference among treatments Duncan's multiple range test (*P* < 0.05)

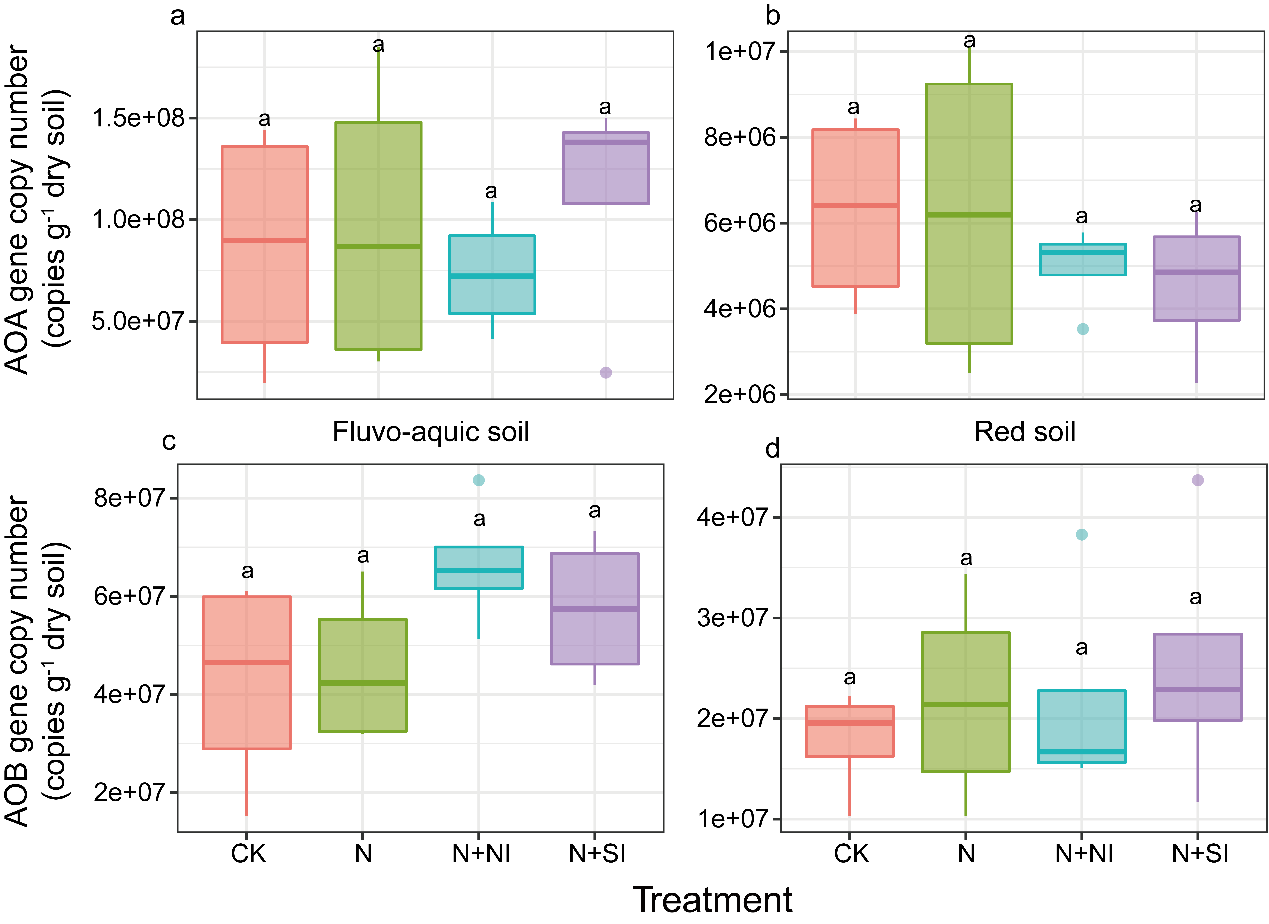

References

Francis, C. A., K. J. Roberts, J. M. Beman, A. E. Santoro, and B. B. Oakley. 2005. Ubiquity and diversity of ammonia-oxidizing archaea in water columns and sediments of the ocean. **102**:14683-14688.

Hallin, S., and P. E. Lindgren. 1999. PCR detection of genes encoding nitrile reductase in denitrifying bacteria. Applied and Environmental Microbiology **65**:1652-1657.

Henry, S., D. Bru, B. Stres, S. Hallet, and L. Philippot. 2006. Quantitative detection of the nosZ gene, encoding nitrous oxide reductase, and comparison of the abundances of 16S rRNA, narG, nirK, and nosZ genes in soils. Applied and Environmental Microbiology **72**:5181-5189.

Rotthauwe, J. H., K. P. Witzel, and W. Liesack. 1997. The ammonia monooxygenase structural gene amoA as a functional marker: molecular fine-scale analysis of natural ammonia-oxidizing populations. Applied and Environmental Microbiology **63**:4704-4712.

Throback, I. N., K. Enwall, A. Jarvis, and S. Hallin. 2004. Reassessing PCR primers targeting nirS, nirK and nosZ genes for community surveys of denitrifying bacteria with DGGE. Fems Microbiology Ecology **49**:401-417.
